# Flexible hidden Markov models for behaviour-dependent habitat selection

**DOI:** 10.1101/2022.11.30.518554

**Authors:** NJ Klappstein, L Thomas, T Michelot

## Abstract

There is strong incentive to model behaviour-dependent habitat selection, as this can help delineate critical habitats for important life processes and reduce bias in model parameters. For this purpose, a two-stage modelling approach is often taken: (i) classify behaviours with a hidden Markov model (HMM), and (ii) fit a step selection function (SSF) to each subset of data. However, this approach does not properly account for the uncertainty in behavioural classification, nor does it allow states to depend on habitat selection. An alternative approach is to estimate both state switching and habitat selection in a single, integrated model called an HMM-SSF. We build on this recent methodological work to make the HMM-SSF approach more efficient and general. We focus on writing the model as an HMM where the observation process is defined by an SSF, such that well-known inferential methods for HMMs can be used directly for parameter estimation and state classification. We extend the model to include covariates on the HMM transition probabilities, allowing for inferences into the temporal and individual-specific drivers of state switching. We demonstrate the method through an illustrative example of African zebra (*Equus quagga*), including state estimation, and simulations to estimate a utilisation distribution. In the zebra analysis, we identified two behavioural states, with clearly distinct patterns of movement and habitat selection (“encamped” and “exploratory”). In particular, although the zebra tended to prefer areas higher in grassland across both behavioural states, this selection was much stronger in the fast, directed exploratory state. We also found a clear diel cycle in behaviour, which indicated that zebras were more likely to be exploring in the morning and encamped in the evening. This method can be used to analyse behaviour-specific habitat selection in a wide range of species and systems. A large suite of statistical extensions and tools developed for HMMs and SSFs can be applied directly to this integrated model, making it a very versatile framework to jointly learn about animal behaviour, habitat selection, and space use.

## 1 Introduction

Wildlife conservation requires an understanding of animal movement and space use (Potts and Börger, 2022). To prioritise areas of conservation interest, it is important to know what habitat features animals use for crucial life processes. These habitat choices (termed ‘habitat selection’) have been studied extensively to understand how animals respond to foraging resources (Bastille-Rousseau et al., 2020), environmental risks (e.g., predators, roads; Fortin et al., 2005; Prokopenko et al., 2017), and other landscape features or resources (e.g., slope, terrain; Fortin et al., 2005; Ellington et al., 2020). Habitat selection can vary between behaviours, which often require different resources (Nicosia et al., 2017). For example, preferred foraging resources may be different from the habitat features that are best suited for other behaviours, such as travelling or resting. Therefore, models for jointly estimating behaviour switching and habitat selection are needed to delineate critical areas for biologically important behaviours.

Step selection functions (SSFs) are a popular framework to jointly model animal movement and habitat selection (Avgar et al., 2016). SSFs assess how animals select habitat by comparing the spatial features (e.g., resources and movement metrics) of selected steps to the surrounding environment (Forester et al., 2009; Avgar et al., 2016). Although SSFs are used to analyse time series data covering long time periods, they usually assume that the selection patterns are constant throughout the movement track. It has been shown that pooling habitat selection parameters over periods that span multiple behavioural states can bias estimates (Roever et al., 2014), but most analyses still ignore this issue. Interactions between movement and habitat covariates can be included to give some insights into how habitat selection varies with movement, but this does not explicitly account for the animal’s behavioural state (Avgar et al., 2016).

Hidden Markov models (HMMs) are commonly used to identify distinct states from animal telemetry data (Morales et al., 2004; Langrock et al., 2012). These states are generally defined based on movement characteristics (e.g., step length or directional persistence), and their dynamics are governed by transition probabilities (e.g., animals may be more likely to stay in their current state than switch). Usually, the estimated states are interpreted as behavioural states, such as resting or travelling (although this will be study-specific; Michelot et al., 2016). It is also common to assess how covariates influence the probability of switching between states (Patterson et al., 2009; Langrock et al., 2012). This is the most common way to incorporate environmental covariates into an HMM (see examples in Morales et al., 2004; Patterson et al., 2009; Michelot et al., 2016), and can further be used to investigate how other temporal (e.g., time of day; Towner et al., 2016) or individual-specific factors (e.g., sex or size; Bacheler et al., 2019) affect animals’ activity state. Although this approach is useful to identify the drivers of behaviours, it falls short of capturing habitat selection directly, as it does not explicitly model individual movement decisions based on habitat features. Therefore, most commonly-used HMMs are inadequate to describe animals’ space use (Glennie et al., 2022).

To assess behaviour-specific habitat selection, some applied studies have employed a two-stage design, in which HMMs are used sequentially with SSFs (e.g., Suraci et al., 2019; Ellington et al., 2020; Clontz et al., 2021; Picardi et al., 2022). In this case, the animal path is segmented into discrete behavioural states using an HMM, and segments of each behaviour are jointly analysed with an SSF to produce state-dependent habitat selection parameters. This two-stage approach is convenient as it uses two widely-used methods, both with user-friendly software and literature (Michelot et al., 2016; McClintock and Michelot, 2018; Signer et al., 2019). However, this approach does not properly account for the uncertainty in behavioural classification, but rather treats the estimated states as data. Further, the state classification is obtained independently of habitat selection. Therefore, the model cannot capture behaviours that are jointly defined by movement and habitat selection, and this may lead to bias or underestimated uncertainty in the habitat selection parameters.

Despite the benefits, it remains complex to analyse habitat selection and behaviour in a unified framework. Nicosia et al. (2017) proposed an integrated model that combines an HMM with an SSF (termed the HMM-SSF, also explored further in Prima et al., 2022). The HMM-SSF accounts for multiple sources of uncertainty and has greater flexibility than a single-stage approach. The SSF component of the model can be used to estimate both movement and habitat selection parameters, and thereby classifies states based on more information than a typical HMM (with only movement covariates). We write this model as a standard HMM with an SSF governing the observation process, and in this paper, we take advantage of this formulation to implement convenient computational methods and extensions. We propose fitting the model using direct numerical maximisation of the likelihood, based on an efficient iterative algorithm called the forward algorithm. We then extend the model of Nicosia et al. (2017) by including covariates on the state transition probabilities, which allows us to estimate effects of external or internal factors on behavioural dynamics. We describe how standard state decoding (i.e., classification into behaviours) can be applied in this context, and show how this spatially-explicit formulation of an HMM can be used to derive space use. Lastly, we present an illustrative analysis of plains zebra (*Equus quagga*) telemetry data as a guide to the application and interpretation of the model.

The HMM-SSF model can viewed in two different ways, and we think that each will appeal to many researchers. It can be viewed as an extension of SSFs, with the addition of behavioural switching, which will be useful to biologists who would like to improve their habitat selection inferences and avoid the pitfalls highlighted by Roever et al. (2014). Alternatively, the HMM-SSF can be described as an extension of the HMMs typically used in animal movement ecology, with the inclusion of habitat selection variables (in addition to movement variables such as step lengths and turning angles). This spatial HMM formulation will be more appropriate in cases where practitioners are interested in capturing animals’ space use (Glennie et al., 2022).

## 2 Methods

### 2.1 Step selection functions

Consider a set of bivariate animal locations {***y***_1_, ***y***_2_, …, ***y***_*T*_ } observed at times 1, 2, …, *T*. An SSF assumes that the probability of an animal taking a given step is determined by both movement constraints and habitat features, such that the likelihood of a step ending at ***y***_*t*+1_ given that it started as ***y***_*t*_ is

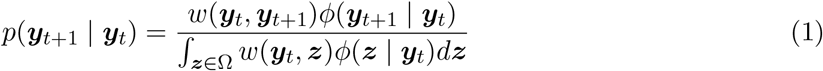

where *w* is a weighting function (evaluated for the step from ***y***_*t*_ to ***y***_*t*+1_) that describes habitat selection in the absence of movement considerations, *ϕ* is a movement kernel that describes how an animal moves in a homogeneous landscape, and Ω is the study area (Forester et al., 2009). Equation 1 can be modified to include higher order dependence (e.g., conditioning on ***y***_*t*− 1_, as described below if turning angle is considered), but this is not always necessary (e.g., Klappstein et al., 2022). The denominator is a normalising constant, which ensures that the SSF is a probability density function with respect to ***y***_*t*+1_ (Potts et al., 2014). Therefore, the entire SSF (which we define as Equation 1, although the nomenclature is inconsistent in the literature) represents the relative attractiveness of the selected endpoint ***y***_*t*+1_ compared to the surrounding habitat.

Although the weighting function *w* and movement kernel *ϕ* can take many forms, defining both as log-linear models allows movement and habitat covariates to be combined into a single selection function (termed an “integrated” SSF; Avgar et al., 2016). That is, the SSF takes the form,

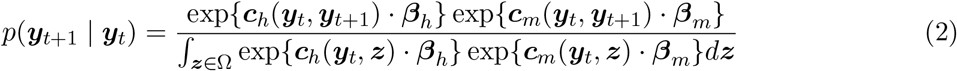

where ***c***_*h*_ and ***c***_*m*_ are vectors of habitat and movement covariates (respectively), ***β***_*h*_ and ***β***_*m*_ are the associated vectors of selection coefficients, and · is the dot product. Through factorisation, this can be simplified to

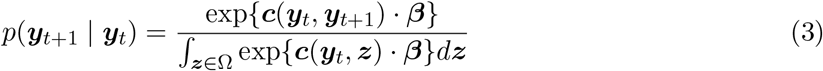

where

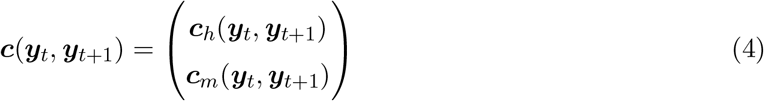

and ***β*** = (***β***_***h***_, ***β***_***m***_). Positive selection coefficients indicate preference for the covariate and negative coefficients indicate avoidance. For a coefficient *β*, exp(*β*) is the multiplicative effect to the SSF of an increase of one unit in the corresponding covariate, all else being constant. This quantity is called the relative selection strength (RSS; Avgar et al., 2017), and is commonly used to interpret habitat selection models. Habitat covariates can be any spatial feature that affects how animals move and use space. There is a wide range of potential covariates, such as foraging resources (e.g., vegetation type, prey density), features that affect the ease of movement (e.g., forest cover, elevation), or proxies of risk (e.g., distance to roads, predator density).

In this model, step lengths can be modelled with various distributions in the exponential family, by including step length (or some transformations) as a covariate in the SSF (Forester et al., 2009; Avgar et al., 2016). In this paper, we model step length with a gamma distribution, which requires including step length and its logarithm as covariates. The general idea is to take the probability density function of ***y***_*t*+1_ | ***y***_*t*_ that would arise from gamma-distributed step lengths, and write it in the form of Equation 3 to identify the relevant covariates ***c***_*m*_. We use a similar idea to model turning angle with a von Mises distribution to capture directional persistence, which requires including the cosine of the turning angle as a covariate. The absolute value of the corresponding selection coefficient is equal to the concentration parameter of the von Mises distribution, either with mean zero (if the selection coefficient is positive) or with mean *π* (if the coefficient is negative). The derivations for step length and turning angle are described in Appendix A, and discussed by Duchesne et al. (2015) and Avgar et al. (2016). Note that the shape and scale parameters of a gamma distribution are stricly positive, whereas the ***β***_*m*_ coefficients are typically estimated with no constraints. As a result, the estimated parameters may sometimes not correspond to a valid step length distribution; in such cases, it may be necessary to use a different distribution, or to constrain the parameters during model fitting.

### 2.2 State-switching SSFs

Following Nicosia et al. (2017), we formulate a state-switching version of an SSF using HMMs. HMMs are doubly stochastic models, in which observation variables {***Y***_1_, ***Y***_2_, …, ***Y***_*T*_ } arise from state-dependent distributions determined at each time *t* ∈ {1, 2, …, *T* } by the latent state variable *S*_*t*_ ∈ {1, 2, …, *K*} (Zucchini et al., 2016). A basic, first-order HMM assumes that the hidden states are a Markov chain, where the state at time *t* is dependent only on the previous state at time *t* − 1. These states are typically considered to represent behavioural states of the animal (e.g., encamped, foraging, travelling), but it is up to the practitioner to draw these inferences based on the estimated state parameters (Morales et al., 2004; Michelot et al., 2016). The state process is characterised by the transition probabilities, given as a *K* × *K* matrix

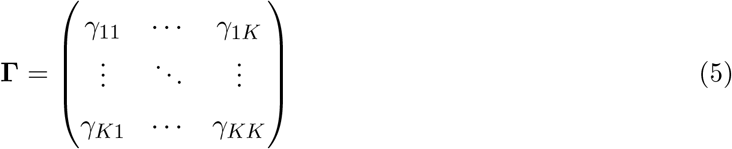

where *γ*_*ij*_ = Pr(*S*_*t*+1_ = *j* | *S*_*t*_ = *i*) is the probability of switching from state *i* to state *j* over one time interval (Langrock et al., 2012). The observation model describes how the data are related to the hidden states. We denote the density of observation ***y***_*t*_ in state *k* as *p*_*k*_(***y***_*t*_) = *p*(***Y***_*t*_ = ***y***_*t*_ | *S*_*t*_ = *k*).

The HMM-SSF is a special case of an HMM, where the distributions *p*_*k*_ are given as SSFs with state-specific parameters. At each time *t*, an animal selects a step according to one of *K* SSFs, as determined by the state at *t*, where each SSF has its own set of selection coefficients (defining movement patterns and habitat preferences in that state). Therefore, the density of a step ending at ***y***_*t*+1_ given that it is in state *k* and started at ***y***_*t*_ is given by the following log-linear SSF,

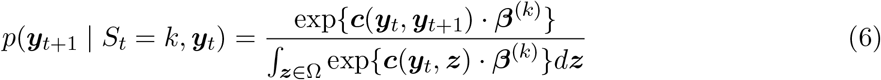

where ***c*** is a vector of covariates and ***β***^(*k*)^ are the associated selection coefficients in state *k*. The likelihood is also conditional on ***y***_*t*− 1_ if turning angle is included, as three successive locations are required for its calculation. The HMM-SSF relaxes the usual assumption of SSF models that successive steps are independent, by inducing dependence through the state process. For example, high probabilities on the diagonal of the transition matrix might imply that a step with high selection for a particular resource is likely to be followed by another step with a similar level of selection. Likewise, it captures some autocorrelation in the speed and directionality of movement, through the movement covariates. This is an improvement over simple step selection functions, where the selection is averaged over all steps. In the rest of this paper, we describe how well-known algorithms and extensions developed for HMMs and SSFs can be adapted to the present context.

#### 2.2.1 Time-varying transition probabilities

We extend the model of Nicosia et al. (2017) to include covariates on the transition probabilities. At time *t*, each transition probability is linked to covariates using a multinomial logit link

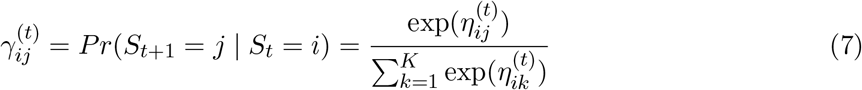

with the linear predictor for *P* covariates 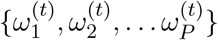 given as

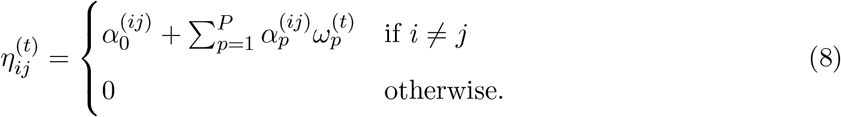

For each transition probability, 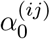 is an intercept parameter, and 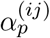 measures the effect of the *p*-th covariate *ω*_*p*_ (Michelot et al., 2016). Note that transition probability covariates will be different to those included in the SSF. Generally, SSF covariates are spatial features that affect step-level movement decisions, whereas covariates included on the transition probabilities are expected to determine the probability of moving into each behavioural state. Among others, transition probability covariates could be temporal variables (e.g., time of day, season) or individual-level attributes (e.g., body condition, sex, age; Patterson et al., 2009; Langrock et al., 2012). These covariates are not “selected” or “avoided” by animals, but they may affect habitat selection behaviour, and this is captured by Equation 8.

In some cases, there might be good reasons to include a covariate either in the SSF (i.e., as ***c***(***y***_*t*_, ***y***_*t*+1_) in Equation 6), or on the transition probabilities (i.e., as 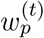 in Equation 8), which would lead to different interpretations. For example, a food resource could be perceived as triggering an animal’s transition into a foraging behavioural state, or the foraging behaviour could be defined as selection for that food resource. Although the covariate could in principle be included in both model components, this might cause estimation problems if the two effects cannot be adequately separated. We suggest using biological knowledge to decide where in the model to include the covariate, and note that this will likely affect the interpretation of the states.

Recently, Prima et al. (2022) also modelled the transition probability of an HMM-SSF as a function of covariates, but did so in a two-stage approach. They defined a binary response based on the probability that the animal transitioned at each time step, which was modelled with a binomial generalised linear model with an environmental predictor. This two-stage approach is not uncommon in HMM analyses (e.g., Breed et al., 2009), but it can be problematic because it does not account for the uncertainty in state classification and it does not allow the state categorisation to depend on environmental covariates. Therefore, direct inclusion of covariates on the transition probabilities is preferable, and relatively straightforward using standard HMM approaches (Patterson et al., 2009; Langrock et al., 2012; Leos-Barajas et al., 2017).

### 2.3 Implementation

#### 2.3.1 Monte Carlo integration

In practice, the integral in the denominator of Equation 6 is analytically intractable (Rhodes et al., 2005). However, we can use Monte Carlo integration (i.e., evaluating the function at random points) to get an approximation of the likelihood (Johnson et al., 2008; Potts et al., 2014). In SSFs, this is often referred to as a “case-control” design, in which random locations (i.e., the controls) are matched spatially and temporally to each observed location (i.e., the case) to approximate each step likelihood. For each time step *t*, we sample *N* control locations {***z***_1*t*_, ***z***_2*t*_, …, ***z***_*Nt*_} and the state-dependent density is approximated as,

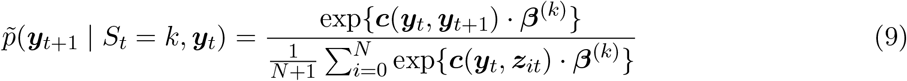

with ***z***_0*t*_ = ***y***_*t*+1_. The term 1*/*(*N* + 1) in the denominator is often omitted when it is constant; here, we include it explicitly as the number of valid control locations may vary between steps due to missing covariate data. In general, the choice of *N* will impact how well the function is approximated, where as *N* →∞ the approximation approaches the true likelihood.

The method to generate control locations in an SSF is an important choice, with potential implications on the precision of the estimation. The simplest method is to randomly simulate controls uniformly over Ω. In practice, it is more common to simulate control locations on a disc that is sufficiently large to encompass the vast majority of the probability mass, as increasing radius size much beyond the maximum observed step does little to improve the estimation (Klappstein et al., 2022). However, uniform sampling can require a large number of random locations to achieve low error, as animals are generally unlikely to take long step lengths and many of the sample controls will have a likelihood close to zero. One way to increase computational efficiency and the precision of the approximation is to preferentially sample where the likelihood is expected to be highest, using importance sampling. If the random locations ***z***_*it*_ are generated from a distribution with probability density function *h* (which typically also depends on the previous observed location ***y***_*t*_), we can re-write the denominator of the approximate likelihood as

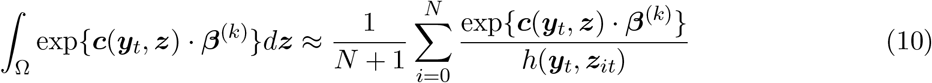

where *z*_0_ = *y*_*t*+1_. For example, random locations could be generated based on gamma-distributed distances from ***y***_*t*_, to take advantage of the animal’s tendency to favour short steps, and *h* would be the corresponding two-dimensional spatial distribution (derived in Appendix A.1; Equation 14).

#### 2.3.2 Direct likelihood maximisation via the forward algorithm

We evaluate the likelihood of the HMM-SSF using a recursive algorithm (i.e., the forward algorithm), which efficiently accounts for all possible state sequences based on the dependence structure of the model (Langrock et al., 2012; Michelot et al., 2016). The model likelihood can be written as

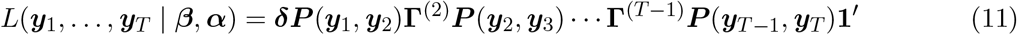

where ***δ*** = (Pr(*S*_1_ = 1), …, Pr(*S*_1_ = *N*)) is the initial distribution of the state process, ***P*** (***y***_*t*_, ***y***_*t*+1_) is a diagonal matrix with *k*-th element 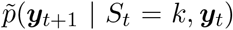 along the diagonal (Equation 9), ***β*** is a vector of the SSF parameters, ***α*** is a vector of the transition probability parameters, and **1**^′^ is a column vector of ones. If there are missing observations, we account for these by defining the corresponding ***P*** as the identity matrix (Langrock et al., 2012). The initial distribution ***δ*** is often susceptible to identifiability issues and, to avoid numerical problems in the estimation, we fix it to be the stationary distribution of the transition probability matrix at time *t* = 1 (Michelot et al., 2016). The Markov chain is not stationary if there are covariates included on the transition probabilities but, even in that case, we assume that the stationary distribution of **Γ**^(1)^ is a good heuristic choice for the initial distribution. To prevent underflow, we implement the forward algorithm for the “scaled” negative log-likelihood of the HMM-SSF, following Zucchini et al. (2016). In the case of multiple individuals in the same data set, we assume that all individuals share the same parameters (“complete pooling”; Langrock et al., 2012; Michelot et al., 2016).

To obtain estimates of all parameters, the negative log-likelihood of the model can then be minimised with respect to ***α*** and ***β*** using a numerical optimiser, and we use optim in R. This approach stands in contrast with the expectation-maximisation (EM) algorithm proposed by Nicosia et al. (2017) to fit the HMM-SSF. Although the two methods will generally converge to the same estimates, it has been argued that direct likelihood maximisation is often faster and easier to implement for HMMs (MacDonald, 2014; Zucchini et al., 2016; Leos-Barajas et al., 2017). Parameter standard errors can then be computed as the square root of the diagonal elements of the negative Hessian matrix Zucchini et al. (2016). For covariate-dependent transition probabilities, confidence intervals are obtained using the delta method (Ver Hoef, 2012). Reliable optimization in HMMs can be sensitive to choice of initial parameter values (Michelot et al., 2016). A common solution is to generate many sets of initial values, use each to fit the model, and finally keep the solution with the lowest negative log-likelihood (implemented in Section 2.4). Another approach could be to use the two-stage approach (HMM then SSF, e.g., Roever et al., 2014; Picardi et al., 2022) to find reasonable starting values, although this would assume that the states are largely defined by movement. We used simulations to check that our model fitting procedure can recover all model parameters when the true data generating process is known. Full details are given in Appendix B.

#### 2.3.3 State decoding

In many situations, it is of interest to estimate the state process *S*_*t*_, a procedure called state decoding. We expect that state decoding will be relevant to most HMM-SSF analyses as a method to identify behavioural phases from movement data. The two main approaches to tackle this problem for HMMs are global and local decoding (Zucchini et al., 2016). Global decoding consists of identifying the state sequence that is most likely to have given rise to the observed data, and can be computed using an efficient iterative algorithm (“Viterbi algorithm”; Zucchini et al., 2016). The output is a sequence of state indices, which can be used to subset or spatially visualise the locations by state. Alternatively, local decoding provides probabilities of occupying each state at each observation time (i.e., the state probabilities), which is often more informative about the uncertainty in state classification (where values close to 0.5 indicates high uncertainty in the state process). Here, we suggest computing the local state probabilities with the forward-backward algorithm (Zucchini et al., 2016). Note that this is equivalent to the approach described by Nicosia et al. (2017) to obtain the state probabilities as a by-product of their expectation-maximisation algorithm. A state sequence can be obtained from local state probabilities, by taking the state with highest probability at each time step. Although this is usually very similar to the Viterbi sequence, it is generally not identical, because they solve different optimisation problems (i.e., global decoding optimises over the full state sequence, rather than at each time step; Zucchini et al., 2016).

#### 2.3.4 Simulating space use

A proposed application of SSFs is to simulate from the fitted model to estimate the utilisation distribution of the animal (Signer et al., 2017), and a similar method can be used for the HMM-SSF. This can give information about areas that are important for different behaviours, which is typically not possible with standard HMM approaches (Glennie et al., 2022). There are several ways to simulate utilisation distributions, which range from generating a single long track (i.e., the steady-state distribution) or several shorter tracks (i.e., a transient distribution; Signer et al., 2017). In this paper, we do not intend to provide guidelines for best simulation practice, but we present an algorithm to simulate data from the HMM-SSF (Appendix B), which can then be used to estimate utilisation distributions. The general steps of the algorithm are to first simulate a state sequence based on the estimated transition probabilities, and then for each time step, simulate the next location from the SSF corresponding to the current state. In practice, this is done by proposing many possible next locations (within a disc), and selecting with probability proportional to their state-specific SSF (Equation 9). Note that the number of proposed end points depends on the size of the disc, and it should be high enough to ensure good sampling coverage, so that bias is not introduced through the simulation. The utilisation distribution can then be estimated using a method such as kernel density estimation on the simulated locations. This can be either an overall distribution based on all locations, or a behaviour-specific distribution if only locations in a given state are kept. The simulations should be designed to best capture the study system, and we provide one example in the next section, with more specific parameters and settings.

### 2.4 Illustrative example

We provide an example to demonstrate the workflow for implementing and interpreting the HMM-SSF. We analysed a track of plains zebra locations collected at a 30-minute resolution from January - April 2014 in Hwange National Park in Zimbabwe (Michelot et al., 2020). The time-series consisted of 7246 observations, with 125 missing locations. We fitted a two-state HMM-SSF, where the SSF component captured selection for a combination of habitat and movement covariates, and the HMM component captured behaviour change as a function of time of day. We included a categorical covariate for vegetation type, with four levels: grassland (reference category), bushed grassland, bushland, and woodland (Figure 1). We modelled step lengths with a gamma distribution (i.e., with step and its log as covariates), and turning angle with a von Mises distribution (i.e., with the cosine of turning angle as a covariate). For transition probabilities, we included time of day *τ* as a cyclic covariate (following Towner et al., 2016), such that the linear predictor in Equation 8 becomes

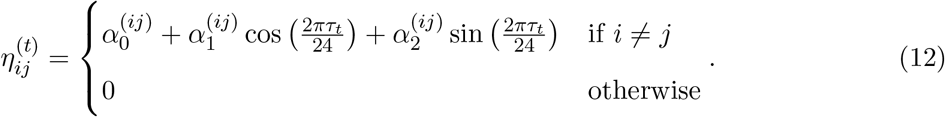

**Figure 1:**
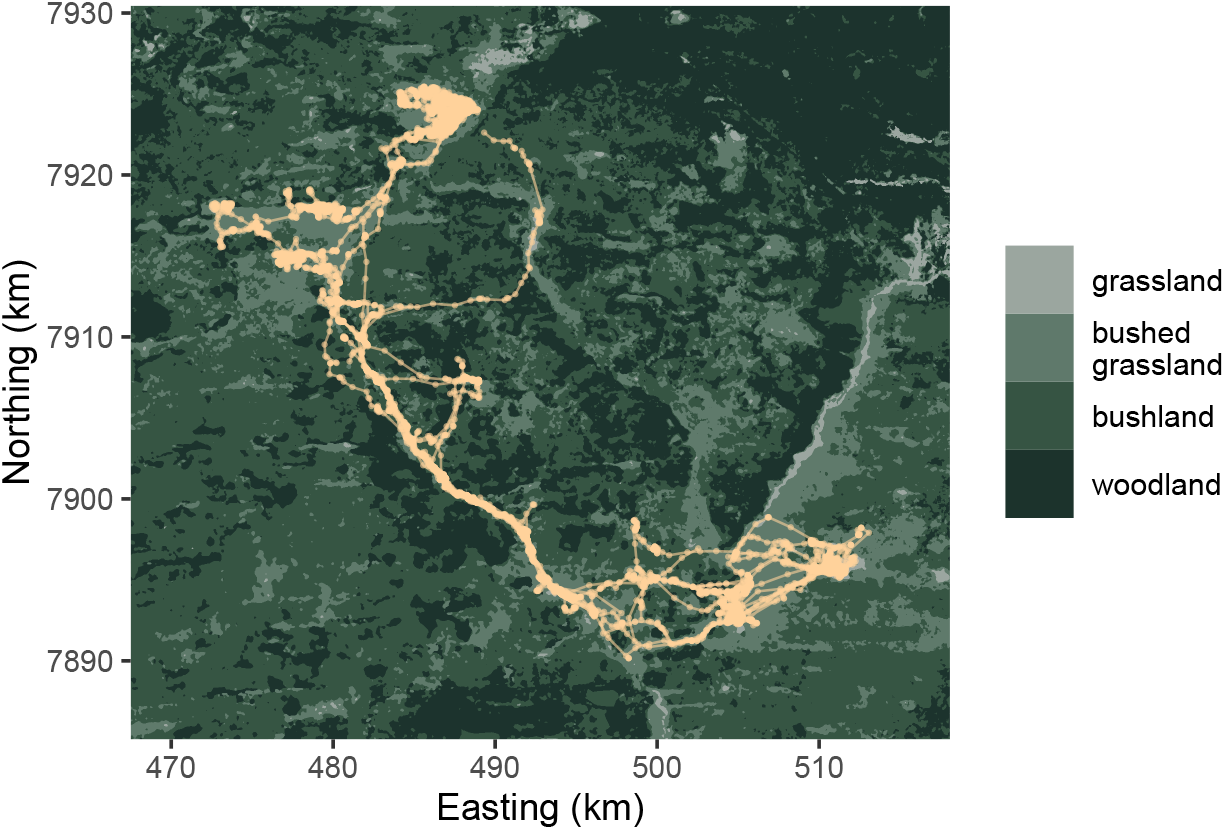
Habitat type map and zebra track (tan). Plot adapted from Michelot et al. (2020).

All fitting was done with the implementation methods described in Section 2.3. To implement the Monte Carlo integration routine, we generated *N* = 25 control locations for each case location. Control step lengths were generated as random draws from a gamma distribution with the mean and standard deviation of the observed data, and turning angles were generated as random draws from a uniform distribution. To ensure convergence of the estimation procedure, we fitted the model with several sets of initial values, spanning different patterns of selection, and chose the model with the lowest negative log likelihood. Lastly, we derived the relative selection strength (RSS) of grassland (i.e., reference category) compared to habitat type *j* as 1*/* exp(*β*_*j*_), where *β*_*j*_ is the selection coefficient for habitat type *j*. The RSS for grassland can be interpreted as how much more likely a zebra is to take a step in grassland, compared to the other habitat type.

We determined the most likely sequence of states using the Viterbi algorithm, and the state probabilities at each time step using the forward-backward algorithm (Michelot et al., 2016; Zucchini et al., 2016). In addition to the transition probabilities, we derived stationary state probabilities as functions of the time of day. For a given time of day, these provide some indication of the probability of being in each state, which is often useful for interpretation (Patterson et al., 2009). In practice, they are derived as the stationary distribution of the transition probability matrix for each time of day. Based on the fitted model, we also simulated a utilisation distribution. We generated five long tracks of 100,000 locations at a 30-minute interval. First, the state sequence was simulated based on the time of day, which was initialised as the first observed time. Then, each location was selected (with probabilities given by their SSF values) from a possible 1000 endpoints on a disc with the radius *r* = max(*L*) × 1.1, where max(*L*) is the maximum observed step length. We chose to run five tracks (rather than one long track), so that we could run each simulation in parallel. The starting location of each track was chosen randomly from the observed data, and we removed the first 1000 locations of each track to reduce the effect of this choice. We used a reflective boundary condition, in which each simulated track was constrained to stay within the study area by fixing the SSF of any point outside the boundaries to zero. We then estimated the utilisation distribution by applying kernel density estimation (bandwidth = 2) to the simulated locations.

## 3 Results

The simulation results (in Appendix B) suggested that all components of the model were generally estimated well, including step length, turning angle, and habitat selection parameters, as well as transition probabilities as functions of a covariate. However, one habitat selection parameter was estimated poorly, which indicated that bias can arise when the spatial scale of covariate autocorrelation is much larger than the scale of movement.

In the zebra analysis, we identified two states with distinct movement and habitat selection patterns. Table 1 provides a full list of selection parameter estimates with uncertainty, but here, we discuss these in terms of the parameters of the state-specific step length distribution (gamma with mean 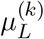 and standard deviation *σ*^(*k*)^; both in km) and von Mises distribution (angular concentration *κ*^(*k*)^ and mean 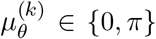). State 1 was identified as a slow state 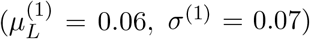 with no directional persistence 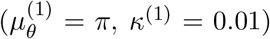. State 2 was characterised by faster movement 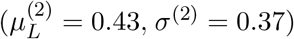 with higher directional persistence 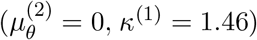 (Figure 2a, 2b). We consider these to be encamped and exploratory behaviours, respectively (Morales et al., 2004).

**Table 1:** Estimated parameters from the HMM-SSF fitted to zebra telemetry data. Note, these are the untransformed β estimates and do not directly represent the mean and variance of the assumed gamma distribution; the turning angle (given by θ) parameter represents the angular concentration of the von Mises distribution. L is the step length (km) and τ is the hour of the day.

|  | State/transition | Covariate |  | Estimate (95% CI) |
| --- | --- | --- | --- | --- |
| **Movement (SSF)** | *Encamped* | step length $L$ | $\beta_1^{(1)}$ | -13.6 (-14.8, -12.3) |
| | | $\log(L)$ | $\beta_2^{(1)}$ | -1.18 (-1.21, -1.15) |
| | | $\cos(\theta)$ | $\beta_3^{(1)}$ | -0.01 (-0.07, 0.04) |
| | *Exploratory* | step length $L$ | $\beta_1^{(2)}$ | -3.11 (-3.36, -2.86) |
| | | $\log(L)$ | $\beta_2^{(2)}$ | -0.65 (-0.78, -0.53) |
| | | $\cos(\theta)$ | $\beta_3^{(2)}$ | 1.46 (1.33, 1.59) |
| **Habitat (SSF)** | *Encamped* | bushed grassland | $\beta_4^{(1)}$ | 0.21 (0.03, 0.39) |
| | | bushland | $\beta_5^{(1)}$ | -0.19 (-0.46, 0.08) |
| | | woodland | $\beta_6^{(1)}$ | -0.56 (-1.08, -0.05) |
| | *Exploratory* | bushed grassland | $\beta_4^{(2)}$ | -1.00 (-1.16, -0.83) |
| | | bushland | $\beta_5^{(2)}$ | -2.19 (-2.42, -1.96) |
| | | woodland | $\beta_6^{(2)}$ | -1.96 (-2.26, -1.66) |
| **Transition probabilities (HMM)** | *Encamped – exploratory* | intercept | $\alpha_0^{(1,2)}$ | -1.83 (-1.98, -1.69) |
| | | $\cos(\frac{2\pi\tau}{24})$ | $\alpha_1^{(1,2)}$ | 0.03 (-0.14, 0.21) |
| | | $\sin(\frac{2\pi\tau}{24})$ | $\alpha_2^{(1,2)}$ | 0.81 (0.63, 0.99) |
| | *Exploratory – encamped* | intercept | $\alpha_0^{(2,1)}$ | -1.26 (-1.40, -1.12) |
| | | $\cos(\frac{2\pi\tau}{24})$ | $\alpha_1^{(2,1)}$ | 0.15 (-0.02, 0.33) |
| | | $\sin(\frac{2\pi\tau}{24})$ | $\alpha_2^{(2,1)}$ | 0.14 (-0.04, 0.32) |

**Figure 2:**
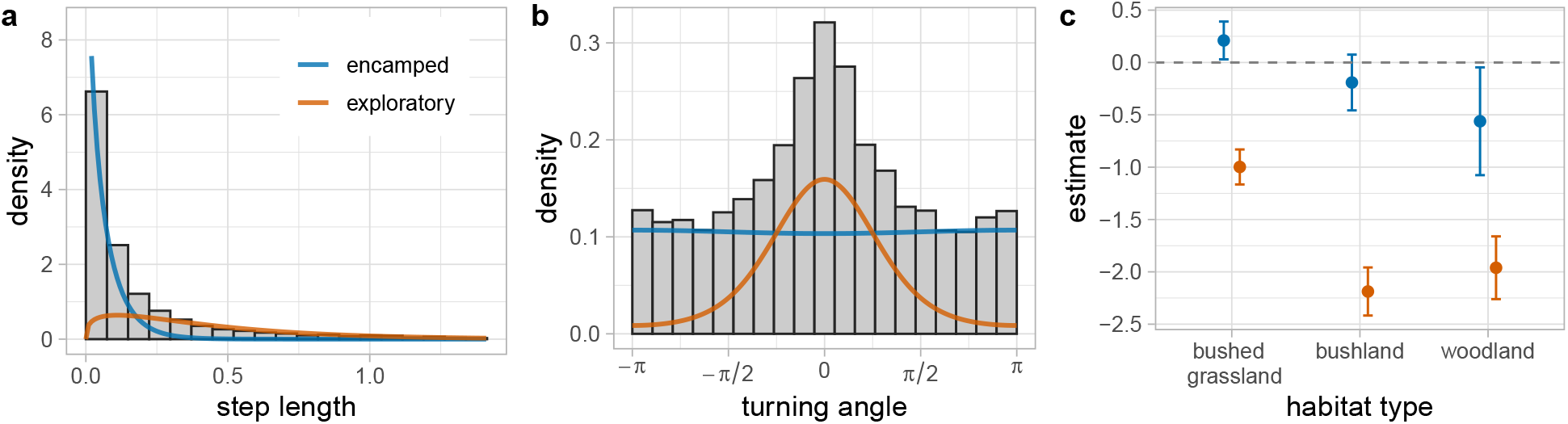
Movement and habitat selection estimates in zebra analysis. Estimated (a) step length and (b) turning angle distributions, weighted by the number of observations predicted to be in each state. The histograms show the empirical distributions of the data. The step length (x-axis) was truncated to the 99th percentile for visualisation purposes. (c) Habitat type parameter estimates (grassland is the reference category, i.e., corresponding to zero).

Habitat selection patterns varied between the two states, although both showed generally high selection for the habitat types with higher grassland cover (Figure 2c). In the encamped state, the zebra selected for grassland over all habitat types, except for bushed grassland, which had a positive selection coefficient. Compared to bushed grassland, the grassland (i.e., reference) RSS was 0.8 (i.e., the zebra was 0.8 times as likely to select grassland than bushed grassland). When encamped, the zebra was more likely to choose grassland over bushland and woodland: the RSS was 1.2 compared to bushland, and 1.8 compared to woodland. Only bushland coefficient had a 95% CI that overlapped zero, but the uncertainty was generally high and the other CIs were also close to overlapping zero (Figure 2c).

Habitat selection was stronger in the exploratory state, where there was clear avoidance of all habitat types relative to grassland and no CIs overlapped zero. The grassland RSS was 2.7, 8.9, and 7.1 for bushed grassland, bushland, and woodland (respectively). Positive selection for grassland is consistent with previous results from Michelot et al. (2020), and may represent selection for their main foraging resources. However, neither state seems to fully capture foraging behaviour in this example, which may explain the selection for grassland or bushed grassland in both states. This may suggest that a 3-state model might be better able to distinguish between biological behaviours in this example (see Pohle et al., 2017, for guidance on the number of states).

We found an effect of time of day on the transition probabilities and on the stationary state probabilities (Figure 3). There was an increase in the probability of transitioning into exploratory in the morning (highest at approximately 07:00), but no strong effect on the probability of transitioning from exploratory to encamped (Figure 3a). The stationary probability of being in the encamped state was highest between 15:00 and 23:00 (peak at roughly 19:00), and lowest between 03:00 and 11:00 (trough at roughly 07:00; Figure 3b). This suggests that this zebra was less active in the late afternoon and evening, and more active in the morning. Maps of the locations classified in each state by the Viterbi algorithm confirm that the encamped state tended to be localised and clustered, where the exploratory state was more spatially diffuse (Figure 4a). The local state probabilities were in agreement with the Viterbi sequence, and the highest state probability was larger than 0.75 for about 85% of time steps (Figure 4b).

**Figure 3:**
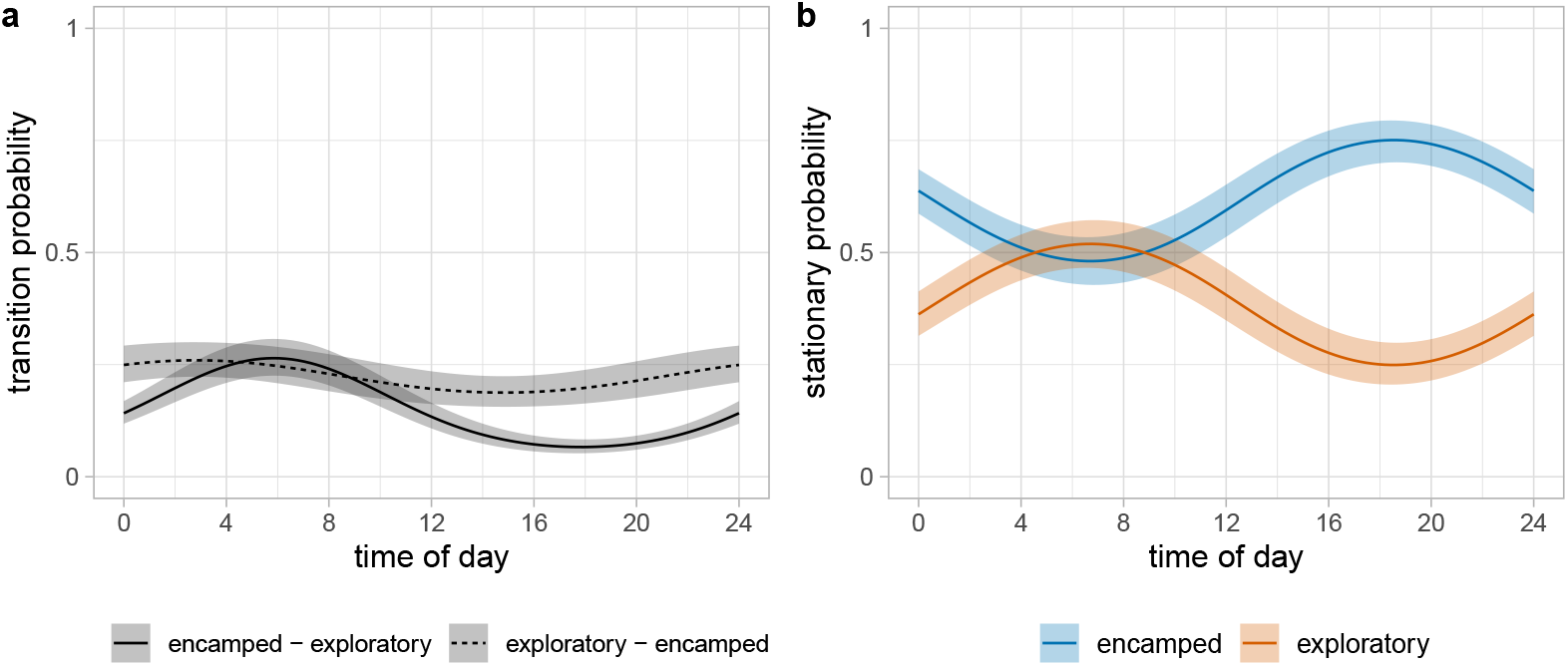
Effect of time of day on transition probabilities in zebra analysis. (a) Estimated transition probabilities γ_12_ and γ_21_ as functions of time of day. (b) Derived stationary state probabilities as functions of time of day. The shaded areas are 95% pointwide confidence bands.

**Figure 4:**
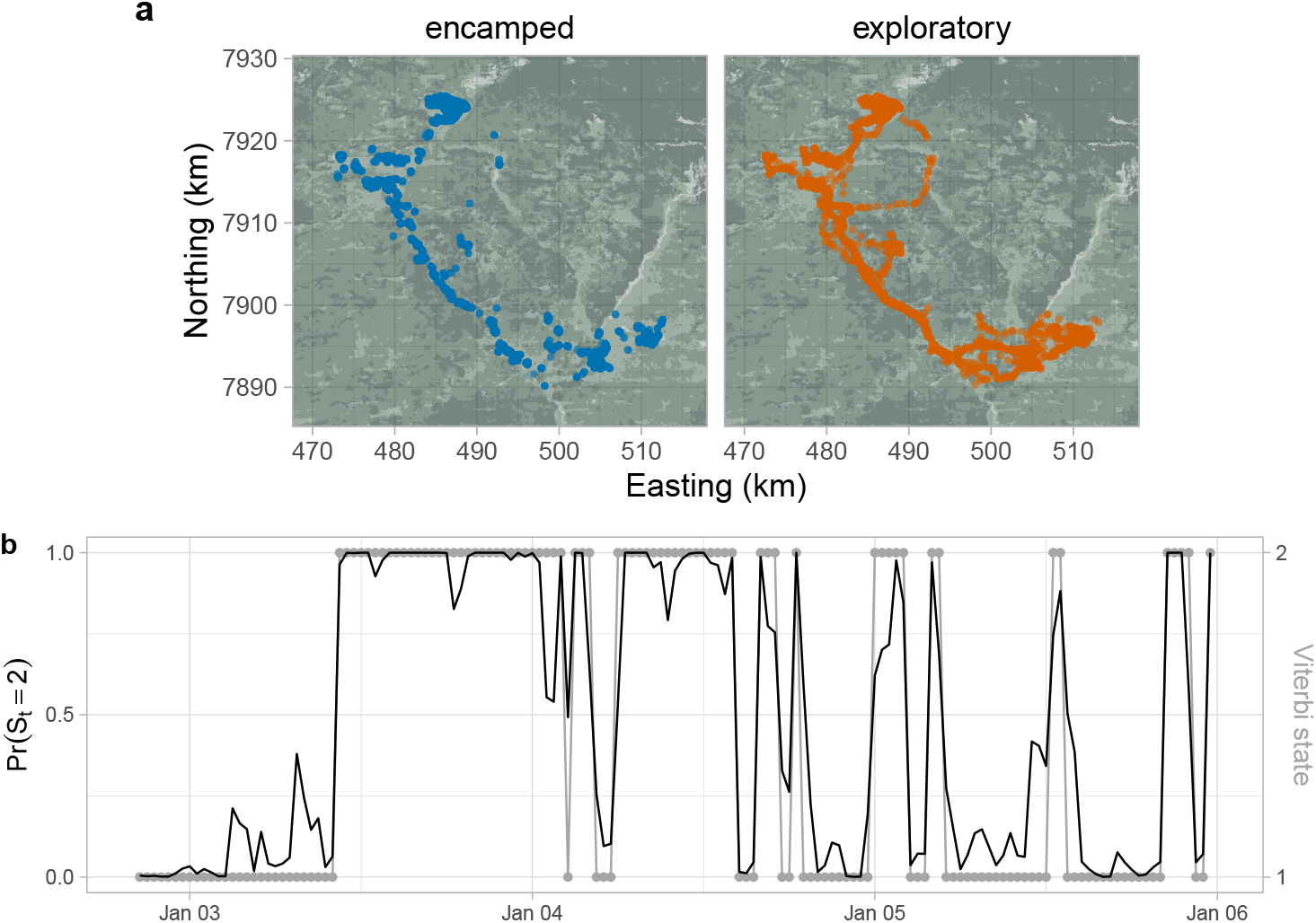
State decoding for zebra example. (a) Locations in each state, estimated via global decoding (Viterbi algorithm). (b) Estimated probability of being in state 2 (black line) and Viterbi sequence (grey points, where 1 is encamped and 2 is exploratory) over a few days chosen for visualisation purposes (note separate y-axes).

We showed that it is possible to estimate large-scale space use from the HMM-SSF, via simulation (Figure 5). The SSF parameter indicated that movement was driven by selection for grassland and bushed grassland, and this was clearly captured in the simulated utilisation distribution.

**Figure 5:**
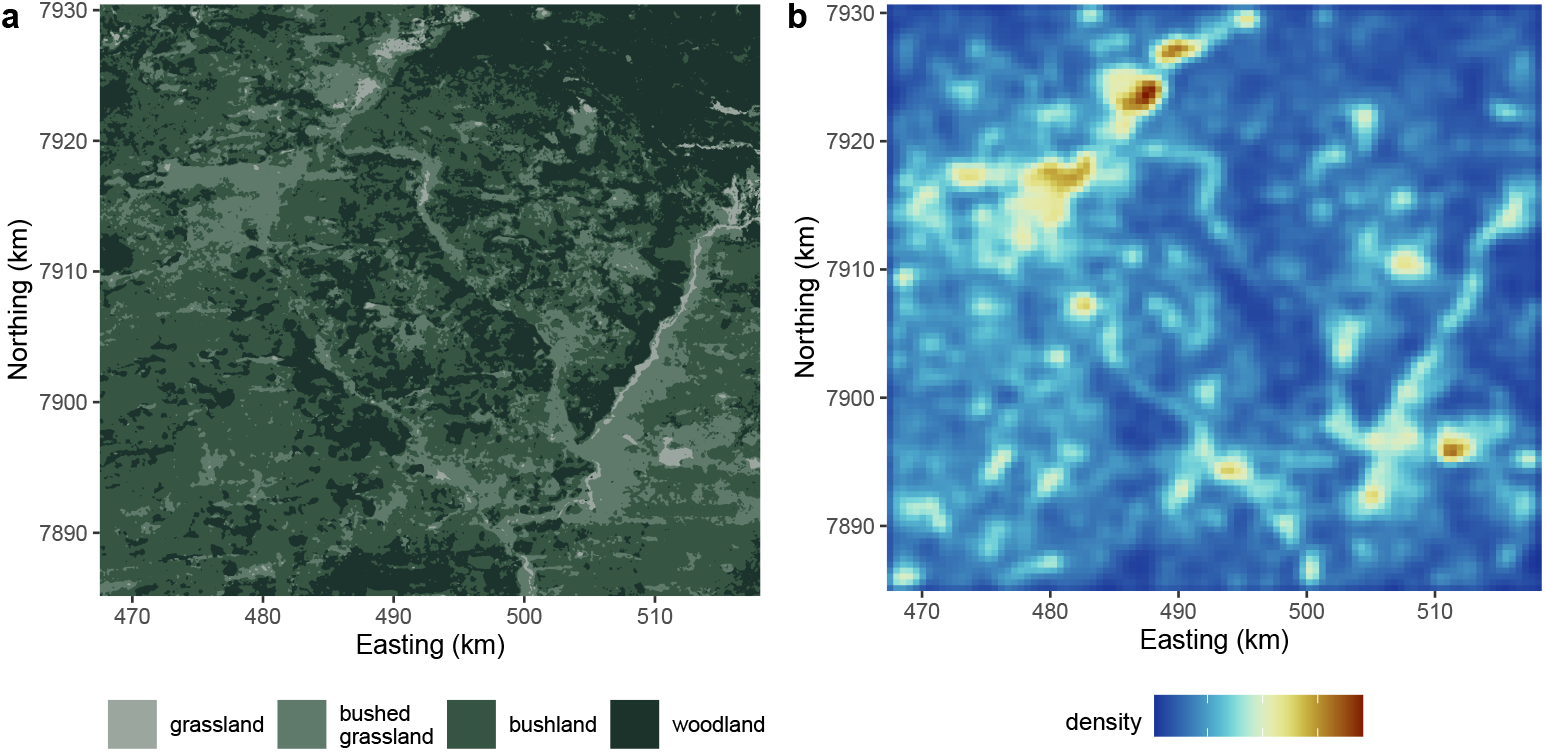
(a) Habitat type map used in the analysis. (b) Estimated utilisation distribution (where blue is lowest density and brown is highest), derived from five simulated tracks of 100,000 locations each. The tracks were simulated based on the estimated HMM-SSF parameters.

## 4 Discussion

In this paper, we focused on developing the flexibility and applied utility of the HMM-SSF, building on recent works by Nicosia et al. (2017) and Prima et al. (2022). Depending on the aims and experience of the practitioner, the HMM-SSF can be viewed as a standard HMM with a habitat selection observation process, or as an SSF that allows for state-switching dynamics.

Viewing this model as a standard HMM opens the way for a wide range of computational tools and extensions, including direct optimisation of the likelihood with the forward algorithm, local and global decoding of the state process, and covariate effects on the transition probabilities. Including additional observation variables in HMMs has been advocated as a method to identify biologically-relevant behavioural states (McClintock et al., 2013), but most HMM analyses still focus solely on movement. In the HMM-SSF, states from are defined by both movement and habitat selection characteristics, and state transitions can also be modelled based on covariates. This approach will be particularly effective when habitat variables are closely linked to behaviour (e.g., a foraging resource will help identify foraging behaviour), and when state transitions depend on temporal or individual-specific factors (e.g., behaviours occur seasonally, or vary by sex, age, etc.).

The HMM-SSF can also be viewed as an improvement over SSFs, via the inclusion of multiple behavioural states. The estimated selection coefficients are interpreted as any other integrated SSF parameters, and can be transformed into relative selection strength and average effect size (following Avgar et al., 2017). The ability to separate behavioural states better accounts for temporal autocorrelation in the data, and can reveal nuanced patterns of habitat selection that would disappear in a simple SSF (e.g., an animal alternating between selection and avoidance of some spatial feature). Deriving space use from SSFs is a commonly desired application, as largescale patterns of interest arise from small-scale movement decisions (Signer et al., 2017; Potts and Börger, 2022). Space use from the HMM-SSF considers non-homogeneous selection parameters, and should be an improvement on single-state SSFs. Although we focused on deriving an overall distribution, there are likely to be many cases where the state-specific distributions are of interest (i.e., when resource selection varies strongly between states), and these are straightforward to derive from the simulated data. We intend our simulation to be a simple illustration, and more refined stochastic and analytical methods could be explored for the HMM-SSF (e.g., methods to upscale from SSFs are reviewed in Potts and Börger, 2022).

The HMM-SSF also inherits some of the limitations of its predecessors, such as scale dependence and implementation challenges. Similarly to HMMs and SSFs, the model is formulated in discrete time, where the estimated parameters scale-dependent and only describe movement, behaviour, and habitat selection at the time interval of the observed movement step (a recognised problem in movement ecology: Michelot and Blackwell, 2019). Additionally, if the SSF is implemented with Monte Carlo integration, it is important to consider how the sampling scheme may affect the precision of the estimation. To improve the approximation, we suggest using importance sampling based on observed step length distribution. State-specific Monte Carlo sampling may be more suitable in cases where the states have very different movement characteristics, but this would likely be very computationally intensive. Lastly, the increased flexibility of the HMM-SSF requires estimating a large number of parameters, which could lead to numerical issues. A large sample of locations may be needed to ensure enough information is available from the data to reliably identify latent behavioural states.

As data sets become larger with technological innovation, we expect the utility of the HMM-SSF to keep increasing. There are many possible extensions to this framework to study complex ecological phenomena, often with minimal changes to the implementation. For example, extended memory dynamics could be incorporated through a higher-order HMM (i.e., where *S*_*t*_ depends on more than just *S*_*t*− 1_; Langrock et al., 2012), or feedback mechanisms could be explored (Langrock et al., 2012, 2014). Additional information about the state or observation process could be incorporated by formulating the model as a semi-supervised HMM (i.e., where states are known a priori; Leos-Barajas et al., 2017), adding additional HMM data streams (such as physiological or accelerometer data; Leos-Barajas et al., 2017), or by including more complex SSF variables (e.g., energetics; Klappstein et al., 2022). Although integrating movement variables in the SSF is convenient, it assumes distributions from the exponential family. This could be relaxed by including movement variables as separate data streams in the HMM, assuming that they are independent of the SSF variables given the state. Generally, this framework could also be used to add a behavioural component in more general SSF formulations, including some recent work on memory (Thompson et al., 2022), movement energetics (Eisaguirre et al., 2020; Klappstein et al., 2022), or movement-habitat interactions (Avgar et al., 2016; Prokopenko et al., 2017). Therefore, a state-switching habitat selection model has many purposes in ecological research, while remaining accessible and interpretable.

## Acknowledgements

We would like to thank Simon Chamaillé-Jammes for sharing the zebra location data used in the illustrative example. We also thank Ron Togunov for comments on an early version of the manuscript, as well as many discussions about behaviour-dependent habitat selection.

## A SSF details

### A.1 Step length distribution

Here, we derive the SSF covariates needed to model step lengths with a gamma distribution. Following Rhodes et al. (2005) and Forester et al. (2009), the two-dimensional spatial distribution of the endpoint ***y*** of a step (given that it started at ***x***) is related to the distribution of the step length *L* = ||***y*** − ***x***|| as follows

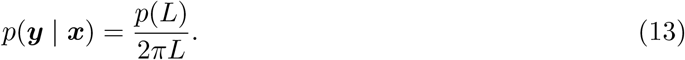

This defines the relationship between the two-dimensional distribution *p*(***y*** | ***x***) and the onedimensional distribution of the distance *L* between the start point ***x*** and end point ***y***. Therefore, the distribution of ***y*** for gamma-distributed step lengths with shape *a* and scale *b* is obtained by replacing *p*(*L*) by the gamma probability density function,

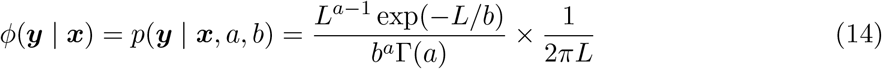

where Γ is the gamma function. To model this in the SSF, we write the movement kernel as the exponential of a linear predictor, such that *ϕ*(***y*** | ***x***) = exp(***c***(***x, y***) · ***β***). Therefore, to identify the terms of linear predictor, we can take the natural log of the movement kernel,

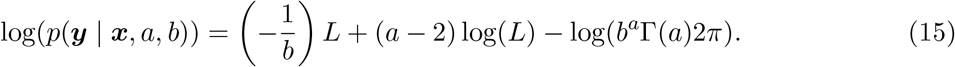

Note that the last term, log(*b*^*a*^Γ(*a*)2*π*), is a constant (i.e., not dependent on the step length) and can be ignored. Finally, this leaves two terms that need to be included in the linear predictor of the SSF: step length *L* and its logarithm log(*L*). The equation gives us the relationship between the gamma distribution parameters and the selection coefficients (here, referred to as *β*_1_ and *β*_2_ for *L* and log(*L*), respectively): *β*_1_ = − 1*/b* and *β*_2_ = *a* − 2. In turn, we can derive mean *μ* and standard deviation *σ* of the step length distribution, in terms of the selection parameters. For the gamma distribution, it is known that

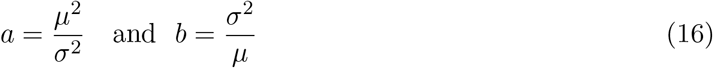

from which we can derive the following

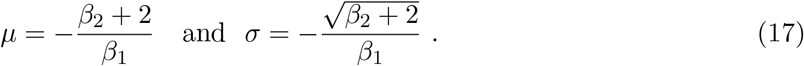

Here, we describe the approach for for the gamma distribution, but this could be applied to other distributions from the exponential family suitable to model step lengths (Forester et al., 2009; Avgar et al., 2016). For example, the exponential distribution is a special case of the gamma distribution, where *a* = 1.

### A.2 Turning angle distribution

Similarly to the procedure used above for step length, we can derive covariates required to model a given distribution of turning angle. Following Avgar et al. (2016) and Nicosia et al. (2017), we focus our attention on the von Mises distribution, with probability density function

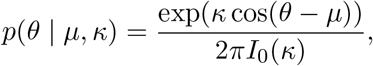

where *θ* ∈ (− *π, π*] is the turning angle, *μ* ∈ (− *π, π*] is the mean parameter, *κ >* 0 is the concentration parameter, and *I*_0_ is the modified Bessel function of the first kind of order 0. Intuitively, *κ* is inversely related to the variance of the distribution, and large *κ* corresponds to a high peak around *μ*.

We first focus on the case where *μ* = 0, corresponding to a tendency to persist in direction, which is the most prevalent in the context of animal movement (but we discuss the case *μ* = *π* below). Assuming *μ* = 0, we take the log of the density function to identify terms that should be included in the SSF,

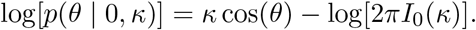

The term log[2*πI*_0_(*κ*)] does not depend on turning angle, and we can therefore omit it. Finally, we get the result that *κ* cos(*θ*) should be added to the SSF linear predictor to model turning angle with a von Mises distribution with mean 0. In practice, this means that the cosine of turning angle is included as a covariate in the model, and the corresponding selection parameter is equal to the concentration parameter of the distribution.

The above assumed that the selection parameter *β*_*θ*_ associated with cos(*θ*) was strictly positive (because *κ >* 0 and *β*_*θ*_ = *κ*). We can relax this assumption by noticing that

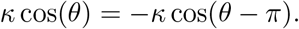

This implies that, if *β*_*θ*_ is negative, we can interpret − *β*_*θ*_ as the concentration parameter of a von Mises distribution centred on *π*. A turning angle distribution centred on *π* corresponds to frequent reversals in direction, which is commonly observed as an artifact of discrete-time observations when animals are inactive.

In summary, including cos(*θ*) as a covariate in the SSF makes it possible to model turning angle with a von Mises distribution, either centred on 0 (if the corresponding selection coefficient *β*_*θ*_ is positive) or centred on *π* (if *β*_*θ*_ is negative). This covers the vast majority of animal movement scenarios, as other values of the mean turning angle are virtually never used.

## B Simulation study

We conducted simulations to verify that our implementation method is able to return the correct parameter values. The basic structure and objective of the simulation study was to: i) simulate a movement track from the HMM-SSF (i.e., with known parameter values), and ii) fit the HMM-SSF with the implementation method described in Section 2.3 and check how well the parameters were estimated.

### B.1 Methods

For all simulations, we followed the same algorithm to produce each movement track:

1. Simulate a state sequence {*S*_1_, *S*_2_, …, *S*_*T*_ }, with *S*_1_ sample with probabilities given by ***δ*** and *S*_2_, …, *S*_*T*_ based on **Γ**
2. Generate starting location *y*_1_ randomly within the study area Ω
3. To generate each *y*_*t*_ in {*y*_2_, *y*_3_, …, *y*_*T*_ }

a. Generate many possible endpoints {*z*_1_, *z*_2_, …, *z*_*J*_ } uniformly on a disc with radius *r*, centred on *y*_*t*_ and evaluate/interpolate relevant covariates.
b. Select *y*_*t*+1_ from the possible endpoints {*z*_1_, *z*_2_, …, *z*_*J*_ } for *j* ∈ 1, 2, …, *J* with probabilities given by the state-dependent SSF (for the known state *S*_*t*_ = *k*), i.e.,

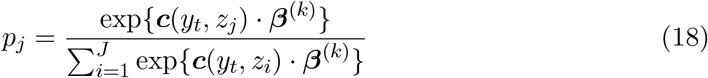

In practice, we generated *J* = 10, 000 endpoints within a disc of radius *r* = 5. *J* was chosen to be very large to reduce the risk of bias arising from the simulation itself. The initial location *y*_1_ was generated from the middle 50% of Ω. Simulations were run for 100 iterations, each for *T* = 750 observations.

We assumed a two-state model meant to represent a generally ideal dataset (i.e., with distinct states and moderate effect sizes), while still being realistic. We assumed that the step lengths followed a gamma distribution (i.e., we included step length and its natural log as covariates) with state-specific means *μ*^(1)^ = 0.5 and *μ*^(2)^ = 2 and standard deviations *σ*^(1)^ = 0.3 and *σ*^(2)^ = 1. We assumed that turning angles followed a von Mises distribution with a mean of 0 (i.e., we included the cosine of the turning angle as a covariate) and state-specific angular concentration parameters *κ*^(1)^ = 0.25 and *κ*^(2)^ = 5. These parameters can be translated to the corresponding distribution and *β* parameters following Appendix A. We also considered selection for two additional habitat covariates, which were simulated using a moving average window with spatial autocorrelation parameter *ρ* (Avgar et al., 2016; Klappstein et al., 2022). Both covariate rasters were simulated with dimensions of 2000 × 2000 with a resolution of 1. The general raster simulation process was to: i) generate a random value for each raster cell from [*U* ∼ (0, 1)], and ii) determine the covariate value based on a circular moving average window with radius *ρ* (i.e., a higher *ρ* indicates more spatial autocorrelation). We defined *ρ*_1_ = 5 for covariate 1 and *ρ*_2_ = 25 for covariate 2 (Figure 6). In state 1, there was selection for covariate 1 (*β* = 3) and selection against covariate 2 (*β* = − 1). This selection pattern was reversed in state 2, with selection against covariate 1 (*β* = − 1) and selection for covariate 2 (*β* = 3).

**Figure 6:**
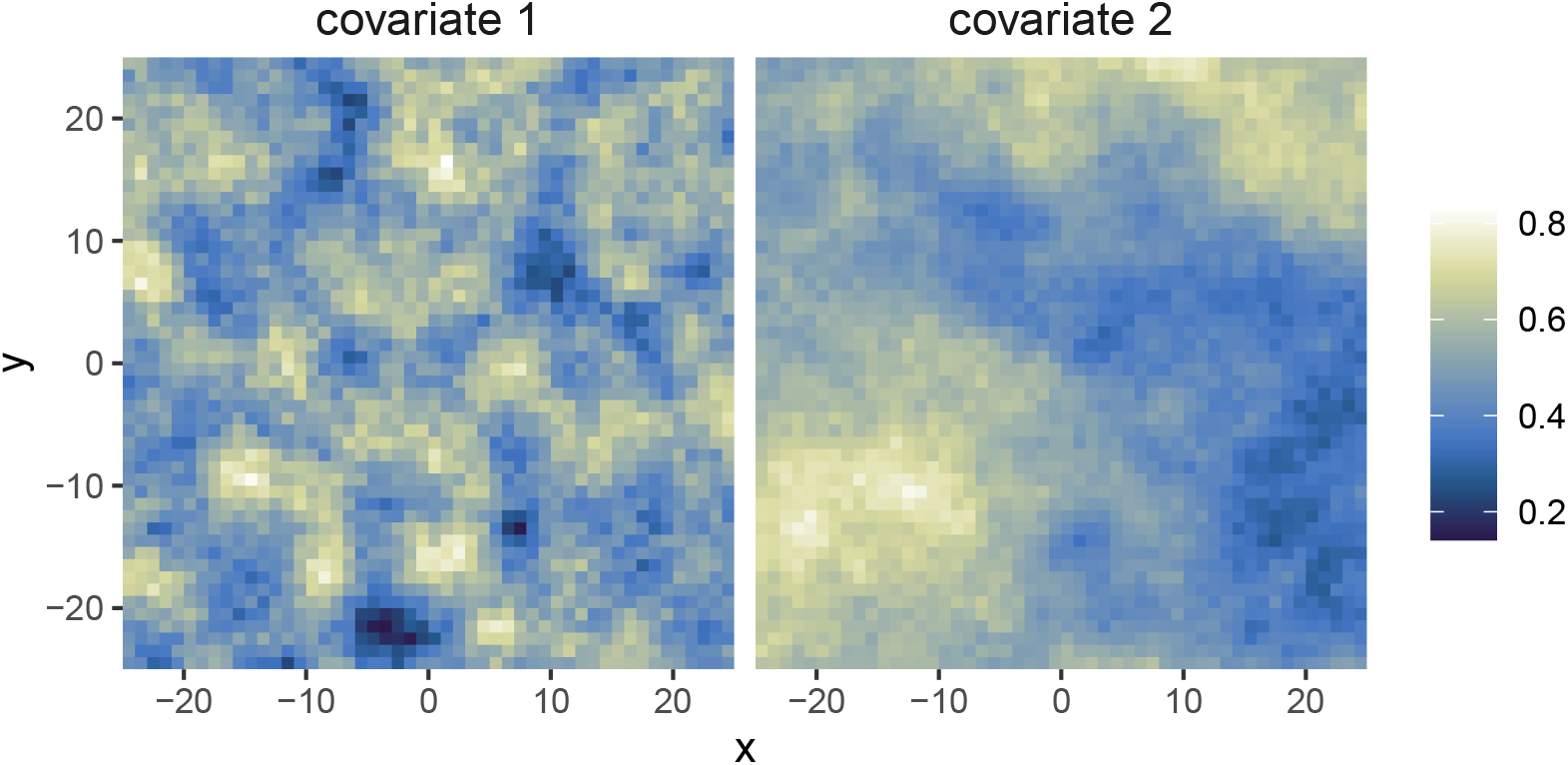
Example of the simulated habitat covariates, with different spatial autocorrelation parameters (ρ_1_ = 5 and ρ_2_ = 25). Shown is the middle cells of the 2000 ×2000 raster, and the color scale indicates the value of the covariate.

We additionally included covariates on the transition probabilities (in addition to the SSF covariates, previously described). We simulated data at a 1-hour time resolution (i.e., 24 steps in a day). Following Towner et al. (2016), we included time of day as a cyclic covariate, such that the linear predictor in Equation 8 becomes

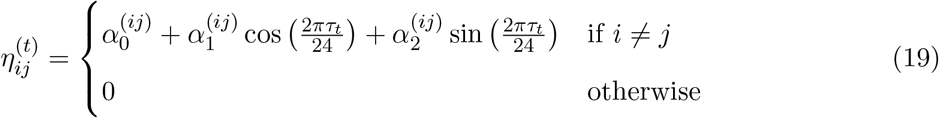

where *τ*_*t*_ is the time of day at time *t*. We set our simulation parameters for the two-state model as,

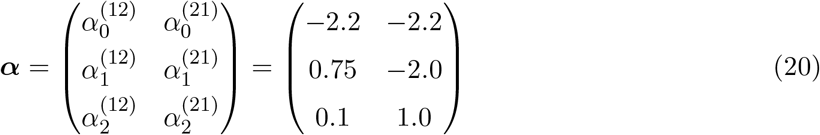

to represent a higher probability of switching into state 1 during the day and a higher probability of switching into state 2 at night. For all simulations, we fitted the model with *N* = 25 gammadistributed control steps.

### B.2 Results

Generally, movement and habitat selection parameters were estimated well, with the exception of the estimate for covariate 2 in state 1 (Figures 7 and 8). This is most likely because there is high spatial autocorrelation in covariate 2 and short step lengths in state 1. This seems to cause bias in the direction of the (positive) selection in state 2 because, when the animal switches into state 1, it will tend to already be in an area of high values of covariate 2, and will not be able to move to low values before switching again. The general pattern of transition probabilities in relation to time of day was also captured, but with higher variance on the transition from state 2 to state 1 (Figure 9). States were decoded correctly 98.2% of the time.

**Figure 7:**
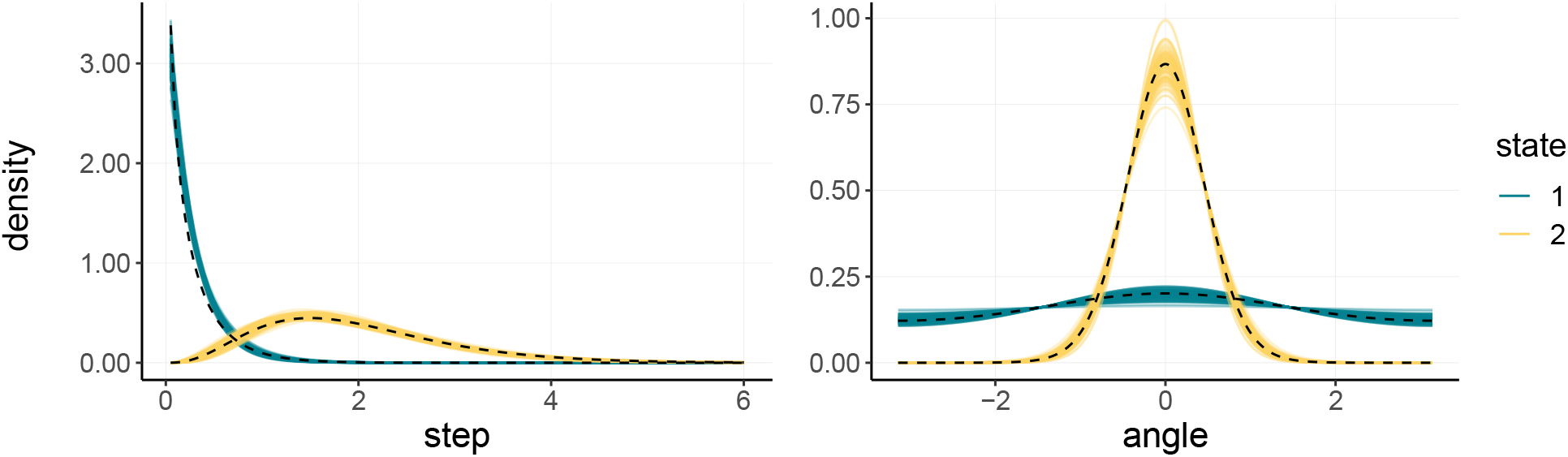
Estimated step length (top) and turning angle (bottom) distributions for all 100 simulation iterations. True distribution shown as the dashed black line.

**Figure 8:**
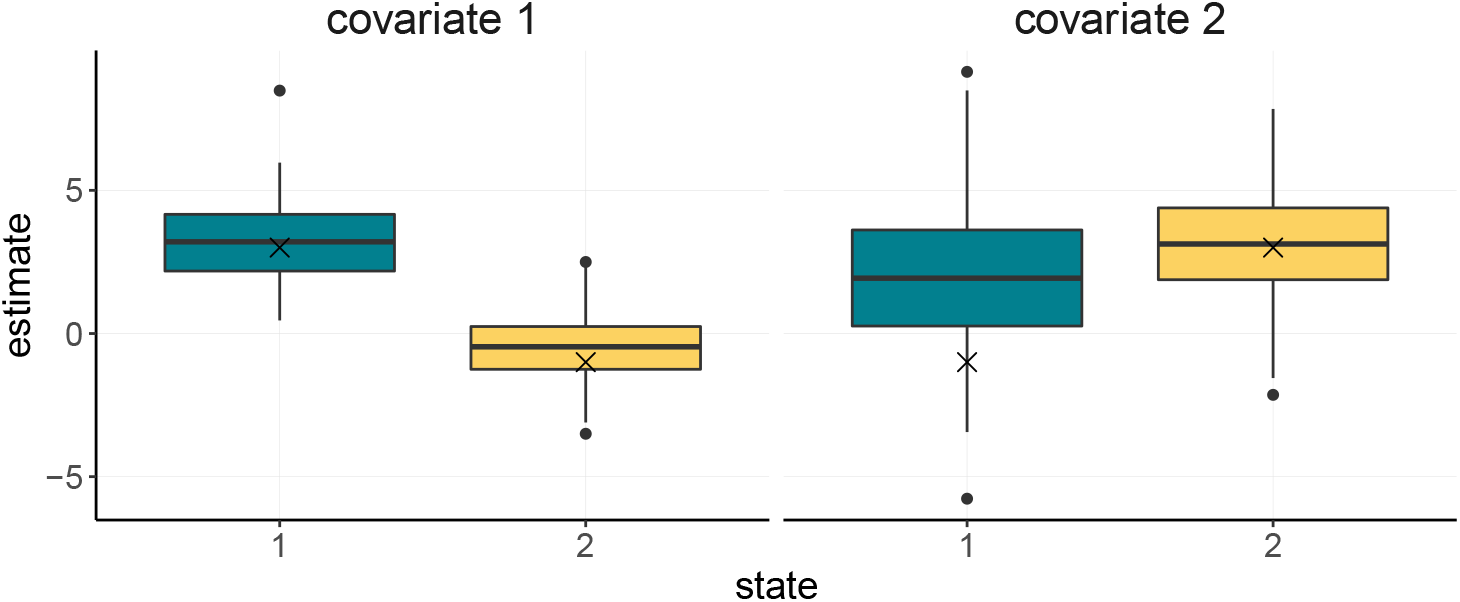
Boxplot of estimated habitat selection parameters (for covariate 1 and covariate 2) for all 100 simulation iterations. True parameter value shown as the black ×, and the middle line is the median of the estimates.

**Figure 9:**
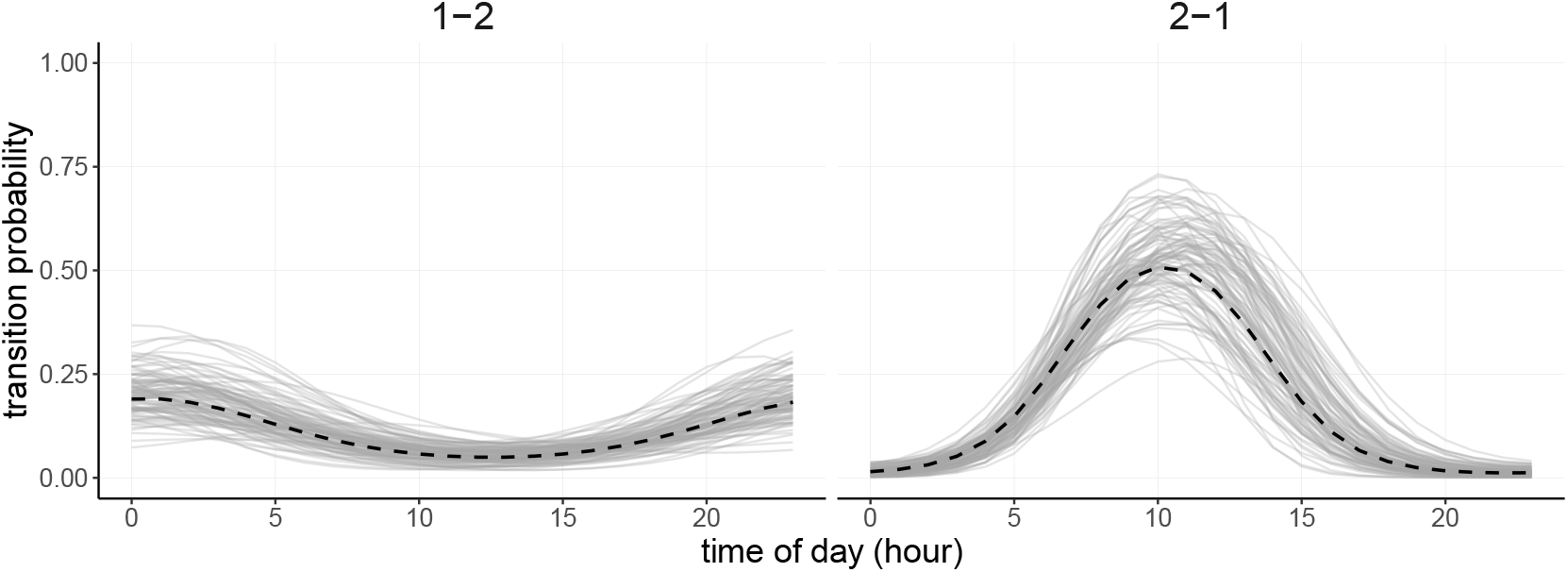
Estimated transition probabilities as a function of time of day, for transitions from state 1 to 2 (1-2) and from state 2 to 1 (2-1). Each line is one of the 100 simulation iteration estimates and the black dashed line is the true relationship.

## References

Avgar, T., Lele, S. R., Keim, J. L., and Boyce, M. S. (2017). Relative Selection Strength: Quantifying effect size in habitat- and step-selection inference. Ecology and Evolution, 7(14):5322–5330.

Avgar, T., Potts, J. R., Lewis, M. A., and Boyce, M. S. (2016). Integrated step selection analysis: Bridging the gap between resource selection and animal movement. Methods in Ecology and Evolution, 7:619–630.

Bacheler, N. M., Michelot, T., Cheshire, R. T., and Shertzer, K. W. (2019). Fine-scale movement patterns and behavioral states of gray triggerfish Balistes capriscus determined from acoustic telemetry and hidden Markov models. Fisheries Research, 215:76–89.

Bastille-Rousseau, G., Wall, J., Douglas-Hamilton, I., Lesowapir, B., Loloju, B., Mwangi, N., and Wittemyer, G. (2020). Landscape-scale habitat response of African elephants shows strong selection for foraging opportunities in a human dominated ecosystem. Ecography, 43:149–160.

Breed, G. A., Jonsen, I. D., Myers, R. A., Don Bowen, W., and Leonard, M. L. (2009). Sex-specific, seasonal foraging tactics of adult grey seals (Halichoerus grypus) revealed by state-space analysis. Ecology, 90(11):3209–3221.

Clontz, L. M., Pepin, K. M., Vercauteren, K. C., and Beasley, J. C. (2021). Behavioral state resource selection in invasive wild pigs in the Southeastern United States. Scientific Reports, 11:6924.

Duchesne, T., Fortin, D., and Rivest, L.-P. (2015). Equivalence between Step Selection Functions and Biased Correlated Random Walks for Statistical Inference on Animal Movement. PLoS ONE, 10(4):e0122947.

Eisaguirre, J. M., Booms, T. L., Barger, C. P., Lewis, S. B., and Breed, G. A. (2020). Novel step selection analyses on energy landscapes reveal how linear features alter migrations of soaring birds. Journal of Animal Ecology, 89:2567–2583.

Ellington, E. H., Muntz, E. M., and Gehrt, S. D. (2020). Seasonal and daily shifts in behavior and resource selection: How a carnivore navigates costly landscapes. Oecologia, 194:87–100.

Forester, J., Kyung Im, H., and Rathouz, P. (2009). Accounting for animal movement in estimation of resource selection functions: Sampling and data analysis. Ecology, 90(12):3554–3565.

Fortin, D., Beyer, H. L., Boyce, M. S., Smith, D. W., Duchesne, T., and Mao, J. S. (2005). Wolves influence elk movements: Behaviour shapes a trophic cascade in Yellowstone National Park. Ecology, 86(5):1320–1330.

Glennie, R., Adam, T., Leos-Barajas, V., Michelot, T., Photopoulou, T., and McClintock, B. T. (2022). Hidden Markov models: Pitfalls and opportunities in ecology. Methods in Ecology and Evolution, 00(n/a):1–14.

Johnson, D. S., London, J. M., Lea, M.-A., and Durban, J. W. (2008). Continous-time correlated random walk model for animal telemetry data. Ecology, 89(5):1208–1215.

Klappstein, N. J., Potts, J. R., Michelot, T., Börger, L., Pilfold, N. W., Lewis, M. A., and Derocher, A. E. (2022). Energy-based step selection analysis: Modelling the energetic drivers of animal movement and habitat use. Journal of Animal Ecology, 91:946–957.

Langrock, R., King, R., Matthiopoulos, J., Thomas, L., Fortin, D., and Morales, J. M. (2012). Flexible and practical modeling of animal telemetry data: Hidden Markov models and extensions. Ecology, 93(11):2336–2342.

Langrock, R., Marques, T. A., Baird, R. W., and Thomas, L. (2014). Modeling the Diving Behavior of Whales: A Latent-Variable Approach with Feedback and Semi-Markovian Components. Journal of Agricultural, Biological and Environmental Statistics, 19:82–100.

Leos-Barajas, V., Photopoulou, T., Langrock, R., Patterson, T. A., Watanabe, Y. Y., Murgatroyd, M., and Papastamatiou, Y. P. (2017). Analysis of animal accelerometer data using hidden Markov models. Methods in Ecology and Evolution, 8:161–173.

MacDonald, I. L. (2014). Numerical Maximisation of Likelihood: A Neglected Alternative to EM? International Statistical Review / Revue Internationale de Statistique, 82(2):296–308.

McClintock, B. T. and Michelot, T. (2018). momentuHMM: R package for generalized hidden Markov models of animal movement. Methods in Ecology and Evolution, 9(6):1518–1530.

McClintock, B. T., Russell, D. J. F., Matthiopoulos, J., and King, R. (2013). Combining individual animal movement and ancillary biotelemetry data to investigate population-level activity budgets. Ecology, 94(4):838–849.

Michelot, T. and Blackwell, P. G. (2019). State-switching continuous-time correlated random walks. Methods in Ecology and Evolution, 10(5):637–649.

Michelot, T., Blackwell, P. G., Chamaillé-Jammes, S., and Matthiopoulos, J. (2020). Inference in MCMC step selection models. Biometrics, 76:438–447.

Michelot, T., Langrock, R., and Patterson, T. A. (2016). moveHMM : An R package for the statistical modelling of animal movement data using hidden Markov models. Methods in Ecology and Evolution, 7:1308–1315.

Morales, J. M., Haydon, D. T., Frair, J., Holsinger, K. E., and Fryxell, J. M. (2004). Extracting more out of relocation data: Building movement models as mixtures of random walks. Ecology, 85(9):2436–2445.

Nicosia, A., Duchesne, T., Rivest, L.-P., and Fortin, D. (2017). A multi-state conditional logistic regression model for the analysis of animal movement. The Annals of Applied Statistics, 11(3):1537–1560.

Patterson, T. A., Basson, M., Bravington, M. V., and Gunn, J. S. (2009). Classifying movement behaviour in relation to environmental conditions using hidden Markov models. Journal of Animal Ecology, 78(6):1113–1123.

Picardi, S., Coates, P., Kolar, J., O’Neil, S., Mathews, S., and Dahlgren, D. (2022). Behavioural state-dependent habitat selection and implications for animal translocations. Journal of Applied Ecology, 59(2):624–635.

Pohle, J., Langrock, R., Van Beest, F. M., and Schmidt, N. M. (2017). Selecting the Number of States in Hidden Markov Models: Pragmatic Solutions Illustrated Using Animal Movement. Journal of Agricultural, Biological, and Environmental Statistics, 22(3):270–293.

Potts, J. R., Bastille-Rousseau, G., Murray, D. L., Schaefer, J. A., and Lewis, M. A. (2014). Predicting local and non-local effects of resources on animal space use using a mechanistic step selection model. Methods in Ecology and Evolution, 5:253–262.

Potts, J. R. and Börger, L. (2022). How to scale up from animal movement decisions to spatiotemporal patterns: An approach via step selection. Journal of Animal Ecology, 00:1–14.

Prima, M.-C., Duchesne, T., Merkle, J. A., Chamaillé-Jammes, S., and Fortin, D. (2022). Multi-mode movement decisions across widely ranging behavioral processes. PLOS ONE, 17(8):e0272538.

Prokopenko, C. M., Boyce, M. S., and Avgar, T. (2017). Characterizing wildlife behavioural responses to roads using integrated step selection analysis. Journal of Applied Ecology, 54:470– 479.

Rhodes, J. R., Mcalpine, C. A., Lunney, D., and Possingham, H. P. (2005). A spatially explicit habitat selection model incorporating home range behavior. Ecology, 86(5):1199–1205.

Roever, C. L., Beyer, H. L., Chase, M. J., and Van Aarde, R. J. (2014). The pitfalls of ignoring behaviour when quantifying habitat selection. Diversity and Distributions, 20(3):322–333.

Signer, J., Fieberg, J., and Avgar, T. (2017). Estimating utilization distributions from fitted stepselection functions. Ecosphere, 8(4):e01771.

Signer, J., Fieberg, J., and Avgar, T. (2019). Animal movement tools (amt): R package for managing tracking data and conducting habitat selection analyses. Ecology and Evolution, 9:880– 890.

Suraci, J. P., Frank, L. G., Oriol-Cotterill, A., Ekwanga, S., Williams, T. M., and Wilmers, C. C. (2019). Behavior-specific habitat selection by African lions may promote their persistence in a human-dominated landscape. Ecology, 0:e02644.

Thompson, P. R., Derocher, A. E., Edwards, M. A., and Lewis, M. A. (2022). Detecting seasonal episodic-like spatio-temporal memory patterns using animal movement modelling. Methods in Ecology and Evolution, 13(1):105–120.

Towner, A. V., Leos-Barajas, V., Langrock, R., Schick, R. S., Smale, M. J., Kaschke, T., Jewell, O. J. D., and Papastamatiou, Y. P. (2016). Sex-specific and individual preferences for hunting strategies in white sharks. Functional Ecology, 30:1397–1407.

Ver Hoef, J. M. (2012). Who Invented the Delta Method? The American Statistician, 66(2):124– 127.

Zucchini, W., MacDonald, I. L., and Langrock, R. (2016). Hidden Markov Models for Time Series: An Introduction Using R. CRC Press, Boca Raton, FL.

